# Neurons need no adaptation to optimally code arbitrarily complex stimuli

**DOI:** 10.1101/2020.05.21.104638

**Authors:** Oren Forkosh

## Abstract

Neural networks seem to be able to handle almost any task they face. This feat involves coping efficiently with different data types, at multiple scales, and with varying statistical properties. Here, we show that this so-called optimal coding can occur at the single-neuron level and does not require adaptation. Differentiator neurons, i.e., neurons that spike whenever there is an increase in the input stimuli, are capable of capturing arbitrary statistics and scale of practically any stimulus they encounter. We show this optimality both analytically and using simulations, which demonstrate how an ideal neuron can handle drastically different probability distributions. While the mechanism we present is an oversimplification of “real” neurons and does not necessarily capture all neuron types, this is also its strength since it can function alongside other neuronal goals such as data manipulation and learning. Depicting the simplicity of neural response to complex stimuli, this result may also indicate a straightforward way to improve current artificial neural networks.

## Main Text

Our brain does a multitude of amazing things. Notably, it has to make sense of the endless stream of complex information flowing from the senses—sight, sound, smell, taste, touch, proprioception, and others—all at once. It is therefore likely that evolution pressure acts to make our neuronal system as efficient as possible (*1*).

Efficiency would mean smaller, more energy-lean brains, with higher processing powers, quicker response rates, increased robustness, and less noise. Yet an optimal neural code is more than just about efficiency—it also about capacity. The brain processes inputs that may span several orders of magnitude in scale or change in their statistics altogether (*2*). The idea that a nervous system can perform ‘optimal coding’ is not just attractive, but was also demonstrated in multiple studies (*3*–*7*).

One way of measuring optimality is by asking how much of the information that the brain receives is lost “on the way” (*8*). In terms of information theory, an optimal system can be described as one which maximizes the mutual information (MI) between what it receives, *x*, and what it yields, *y*. Mutual information is simply a measure of the reduction in entropy, or uncertainty, of a neuron’s output, given the input it received:

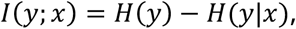

where *I*(*y*; *x*) is the MI between *x* and *y*, *H*(*y*) is the entropy of *y*, and *H*(*y*|*x*) is the conditional entropy, or the remaining uncertainty about *y* once *x* is given. If a neuron encodes the input precisely, the mutual information equals the entropy of the input; otherwise, the MI will have some lower, positive value. If the neuron is not responding to the stimuli at all or producing pure noise, the mutual information declines to zero.

It is typically difficult to use MI to compute the optimal way to encode a signal, but the application of two simple and well-known assumptions leads to a closed-form solution (*9*). The assumptions are that 1. neurons have a limited firing rate, between zero and some upper bound *f*_*max*_ and that 2. neurons tend to increase their spike-rate as the stimuli increase in amplitude. The second assumption is known as Adrian’s law (*2*), however, a similar optimality rule follows if neurons were strictly decreasing their firing rate in response to an increase in the input stimuli (such as retinal OFF cells (*10*)). While these assumptions are an oversimplification of actual neurons it does not mean that deviating from them nullifies our results. Neurons have additional tasks to information transfer (such as data manipulation), but incorporating them in the model makes an analytical analysis difficult.

Taking these two assumptions into consideration, we end up with the following optima transfer function *F*_*opt*_(***x***), relating the stimuli to the spike-rate of the neuron (see methods):

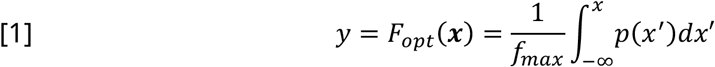

This result shows that an optimal transfer function depends on the probability distribution *p*(*x*) of the input stimuli; or, more precisely, the optimal response is simply the cumulative distribution function of the input stimuli. At first glance at least, this may suggest that optimality requires knowing the statistical properties of each new stimuli (see examples in Figure 1). It turns out, however, that many neurons achieve this optimality without any need for memorizing the statistics of the input.

**Figure 1.**
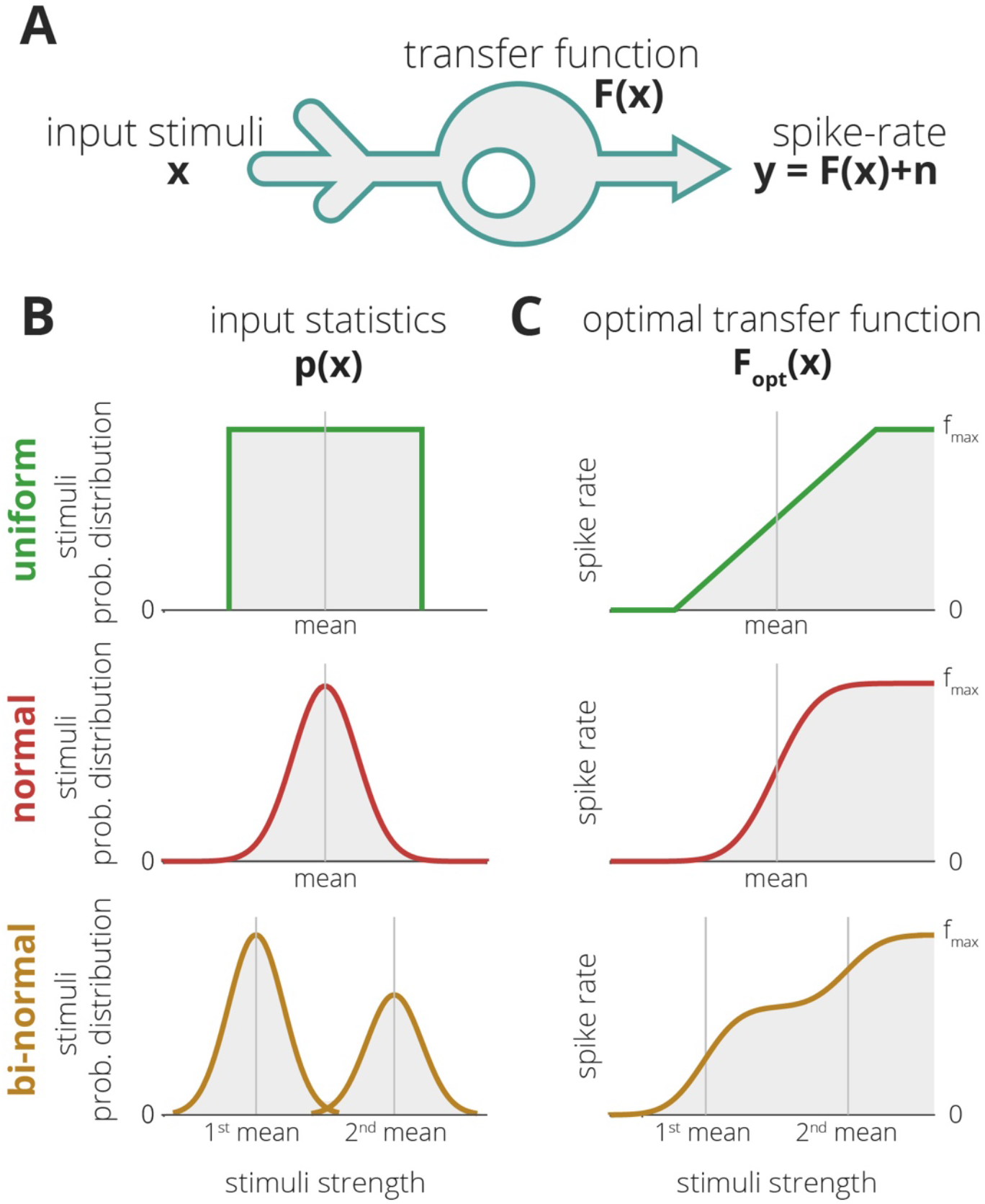
Optimal coding in a single neuron. A. Neurons are small processing units that receive input stimuli x, which modulate their membrane potential to produce all-or-none responses, known as spikes. The number of spikes that a neuron produces in a given time window (which is proportional to the probability of spiking) is referred to as spike rate. The relation between the input stimuli and the spike rate is the neuron's transfer function, and may depend on various factors, such as its geometry, the types of ion-channels it expresses and their distribution, etc. B and C. The optimal transfer function is often defined as one that maximizes the mutual information between the coming signal and the spike-rate of the neuron. In this case, the optimal transfer function equals the cumulative distribution function (CDF) of the input stimuli. We show here three examples (see methods).

Neurons tend to be novelty detectors; that is, they have a stronger response to fluctuations in stimuli (*11*, *12*). In other words, neurons can be seen as differentiators and, turns out, this is also what makes them optimal coders.

The simplest differentiator neuron can be defined by:

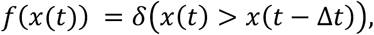

i.e. it produces a spike (denoted here by the delta function) whenever the stimuli *x*(*t*) increase in magnitude within a short timeframe, Δ*t*. In this case, the probability of the neuron to produce a spike at time t (which is proportional to its spike rate, *F*(***x***)) is equal to the probability that *x*(*t*) is larger than *x*(*t* − Δ*t*). In mathematical terms,

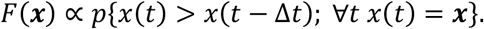

This probability *p*{…} can be computed precisely and equals

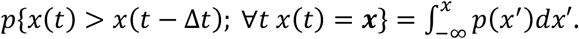

We end up with a spike rate *F*(***x***) that is proportional to

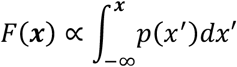

that is, it is exactly proportional to the optimal coder neuron we found according to equation [1] (Figure 2).

**Figure 2.**
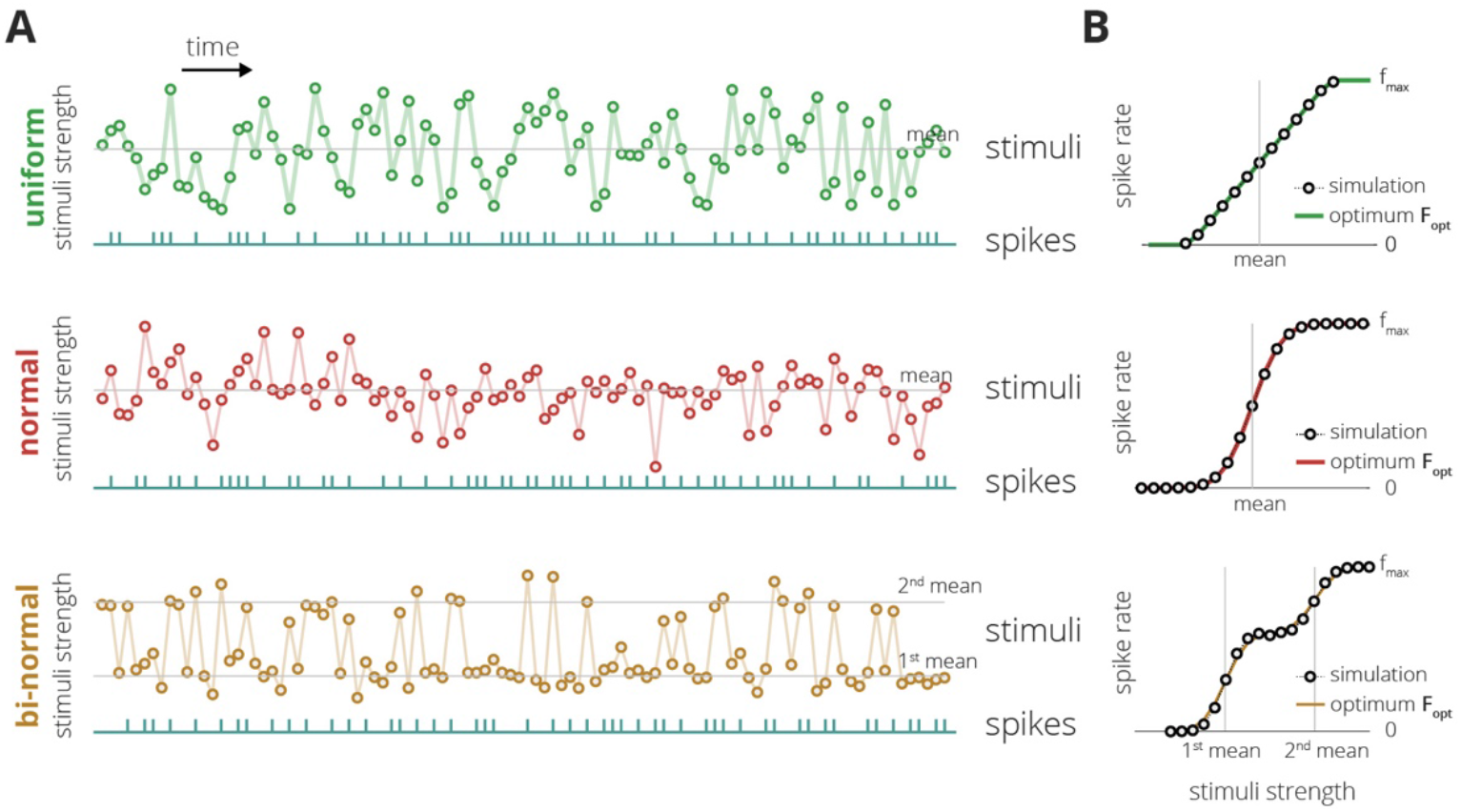
Simple differentiator neurons are optimal coders. A. Random stimuli drawn from the distributions in Figure 1B (100 samples each) and the spiking response of a simple differentiator neuron shown under each sequence (in cyan). Differentiator neurons produce a spike in response to an increase in stimuli amplitude. B. The probability of producing a spike, given the stimuli amplitude (in circles), compared to the optimal model from Figure 1C.

## Discussions

Thus, simple differentiator neurons can drive optimal spike-rate responses to stimuli of different statistics. The model we have shown here is very simplistic and does not take into account factors such as noise, differences between neurons and between brain regions, or varying timescales. Yet the fact that such a simple mechanism can allow neurons to respond efficiently to arbitrary probability functions is surprising. It also provides a basic template for neurons, which allow us to examine neurons in comparison to this ideal representation of a sensory neuron.

To keep this work brief, we only addressed how information is processed by single neurons, yet scaling this idea to a network is straightforward. By introducing noise to the model (*6*) we get variability in the responses of the neurons, especially if this noisiness is reflected in the timing of the spikes we get decorrelated responses to the stimuli without impairing the capacity of each neuron.

The simplicity of this model has another attractive benefit: Research on artificial neural networks (ANN) has shown that applying even the most fundamental properties of actual neurons was enough to revolutionize the field of artificial intelligence. Yet, ANNs, in general, do not make use of the temporal properties of the inputs they receive (apart from at the network level, as in recurrent networks, such as LSTMs (*13*)). This work suggests that we might need to change that, but this change comes at a small cost in terms of model complexity.

## Materials and Methods

Optimal Coding. We assume here that neurons use rate coding to encode their stimuli. The transfer function, which is the relation between the incoming stimuli x and the spike rate y, can be defined as

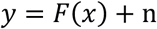

where *n* accounts for noise which, for simplicity, we assume is additive (*14*).

We use a common approach for quantifying the optimality of a neural code, which is measuring the amount of information that a neuron preserves about its stimuli:

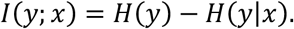

Thus, an optimal code maximizes the mutual information between the inputs x and outputs y, or

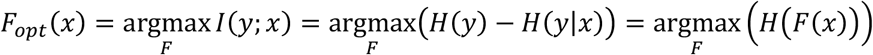

where we can ignore the noise term *n* since it is independent of *F*. This result shows that the only term we need to maximize is the entropy of the neuron’s spike rate.

Assuming that neurons tend to increase their spike rate as the stimuli increases (Adrian’s law (*2*)), we can formulate the dependency of the transfer function on the distribution of the input stimuli (*15*)

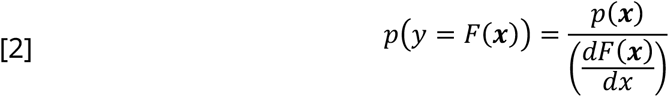

For a bounded random variable, the maximum entropy is achieved for a variable with a uniform distribution. Since a neuron’s spike-rate has an upper bound, *f*_*max*_, an optimal code would, therefore, have a fixed spike-rate, such that

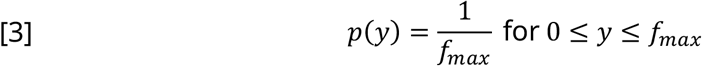

Combining equations [2] and [3], we can derive an exact expression of the optimal response to a given stimulus:

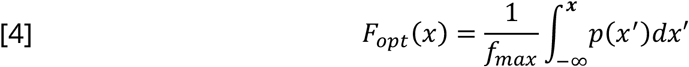

Simulations. To test and illustrate our theory, we used sequences that were randomly sampled from three distributions:

1. Uniform *x*~*U*(−1,1) or 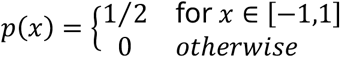
2. Normal *x*~*N*(*μ* = 0, *σ*^2^ = 1) or 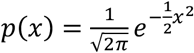
3. Bi-normal distribution with *x*~.6 · *N*(*μ* = 0, *σ*^2^ = 1.8) + .4 · *N*(*μ* = 9, *σ*^2^ = 1.8), that is 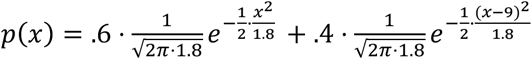

We ran a model differentiator neuron on each sequence, which spiked whenever the stimuli increase in magnitude (i.e., when *x*(*t*) > *x*(*t* − 1)). As the stimuli are continuous, in order to compute the spike rate in Figure 2B, we divided them into n equal bins, *b*_1_, … , *b*_*n*_ (we used n=20). For each bin *b*_*i*_, we computed the probability of a spike for the samples assigned to that bin:

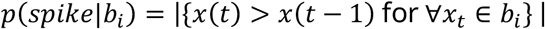

and compared it to the optimal spike-rate in eq. [4] (Figure 2B).

The code used for the simulations is available in https://github.com/OrenForkosh/OptimalCoding

